# The NanoLuc Assay System for Accurate Real-Time Monitoring of Mitochondrial Protein Import within Intact Mammalian Cells

**DOI:** 10.1101/2022.09.23.509159

**Authors:** Hope I Needs, Gonçalo C. Pereira, Jeremy M Henley, Ian Collinson

**Affiliations:** School of Biochemistry, University of Bristol, Bristol BS8 1TD, UK

## Abstract

Only a few proteins (13 in humans) are encoded by the mammalian mitochondrial genome. Therefore, the other mitochondrial resident proteins (>1000) must be recruited *via* specialised import pathways. Protein import is critical for mitochondrial biogenesis and bioenergetic function and health; loss of function has been implicated with a wide range of pathologies. Despite this, our understanding of the kinetic and dynamics of import is somewhat limited, particularly within mammalian cells. Here, we report an adaptation of an assay system, established previously to monitor mitochondrial import into isolated yeast mitochondria, to quantitatively monitor mitochondrial import inside mammalian cells. The reporting is based on a split luciferase, whereby the large fragment is segregated in the mitochondrial matrix and the small complementary fragment is fused to the C-terminus of a recombinant precursor protein destined for import. Following import successively through the TOM complex of the outer membrane and the TIM23 complex of the inner membrane, the complementary fragments combine to form an active luciferase. The resultant luminescent signal provides a sensitive, accurate, free of noise and continuous measure of protein import, enabling mathematical model fitting to identify and understand the steps that make up import. This advance allows detailed mechanistic examination of the transport process in live cells. In addition, the assay will enable characterisation of the protein import when the machinery is challenged; for example, in situations associated with disease. Moreover, the assay is compatible with high throughput for large data set collection and kinetic modelling, as well as for drug screening and characterisation. Our set-up also has the potential to be adapted for the analysis of alternative transport systems and different cell types, and even for multicellular model organisms.

## Introduction

Since most proteins required for mitochondrial function are encoded in the nucleus and synthesised by cytoplasmic ribosomes, efficient targeting and translocation of proteins into mitochondria is critical [1, 2]. In recent years, impaired mitochondrial protein import has been implicated in a range of pathologies, including myopathies and neurodegeneration [3–7]. Therefore, an understanding of the molecular basis for these processes, and how they break down will be crucial for our understanding and alleviation of associated disease.

Until recently, monitoring protein translocation has only been possible by carrying out end-point time course assays *in vitro* in isolated mitochondria, wherein import is detected by radioactivity and Western blotting-based outputs [8–10]. These classical assays have been crucial for seminal studies of our understanding of protein translocation in mitochondria (and bacteria); however, they are laborious and time consuming, and the resultant data are noisy, discontinuous, and not amenable to kinetic analysis and modelling. Moreover, the poor time resolution means that the initial fast phase of the reaction cannot be observed. Another caveat is that the results generated *in vitro* are not necessarily representative of what is happening within the whole cell. Therefore, new approaches are required in order describe the individual steps that make up import and to understand the process *in vivo*.

Several alternative methods to achieve real-time monitoring of protein interactions and trafficking have been described over the past couple of decades, but all have significant limitations, as discussed previously [11, 12]. Split-fluorophore systems [13] are reliant on slow chromophore maturation (slower than the import reaction) which obscures the kinetics of the reaction of interest. Similarly, β-galactosidase assays have been used to monitor protein translocation through the nuclear envelope and plasma membrane [14], and could in theory be applied to mitochondria. However, these assays rely on reporter oligomerisation and product accumulation, hindering its ability to monitor import kinetics. Technology such as SNAP-tags can improve the time resolution of standard Western blotting-based import assays [15], but these are still laborious and cannot generate continuous import data. Thus, there has been a real need for innovative rapid and efficient experimental systems for real-time monitoring of mitochondrial protein import into isolated mitochondria, as well as within live cultured cell systems and *in vivo* within live multicellular model organisms.

NanoLuc Binary Technology (NanoBiT) is a highly sensitive, specific, and stable bioluminescence-based reporter system that utilises a small luciferase subunit derived from a deep-sea shrimp [16]. This technology was originally developed by Promega [11] to monitor protein-protein interactions within living cells. It is a split luciferase, complementation-based system that relies on the binding of 11S (also known as LgBiT), an 18 kDa fragment of NanoLuc, and pep114 (also known as SmBiT), a small, 1.3 kDa peptide of 11 amino acids, which is the final β-strand of NanoLuc [11]. The resulting fully functional NanoLuc enzyme can convert furimazine to furimamide, producing a bioluminescent signal that can be detected with a luminometer [11, 17]. Pep86 (also known as HiBiT) is a high-affinity variant of pep114 (Kd 0.7 nM *versus* 190 μM, respectively), also developed by Promega [11].

Recently, we have adapted and optimised this technology to monitor protein transport across biological membranes [12]. In essence, the 11S fragment is contained within a reconstituted or native vesicle/ organelle/ cell and the small fragment (*e.g*., pep86) is fused to the protein destined for transport – *e.g*., at the C-terminus of a bacterial pre-secretory protein or mitochondrial precursor (Fig. 1A). Protein translocation is then monitored as the pep86-tagged protein crosses the membrane and associates with the encapsulated 11S. Under the conditions deployed the binding of the two fragments to form the active luciferase is an order of magnitude faster than protein transport [12]. Thus, the luciferase signal reports faithfully on the detailed time-resolved kinetics of import. Importantly, it reduces the timescale of data acquisition from days (for classical assays involving radioactivity and/ or Western blotting) to seconds. Its application to the analysis of protein translocation for bacterial secretion has been instrumental in the elucidation of the underlying molecular mechanism for transport [18, 19].

**Fig. 1.**
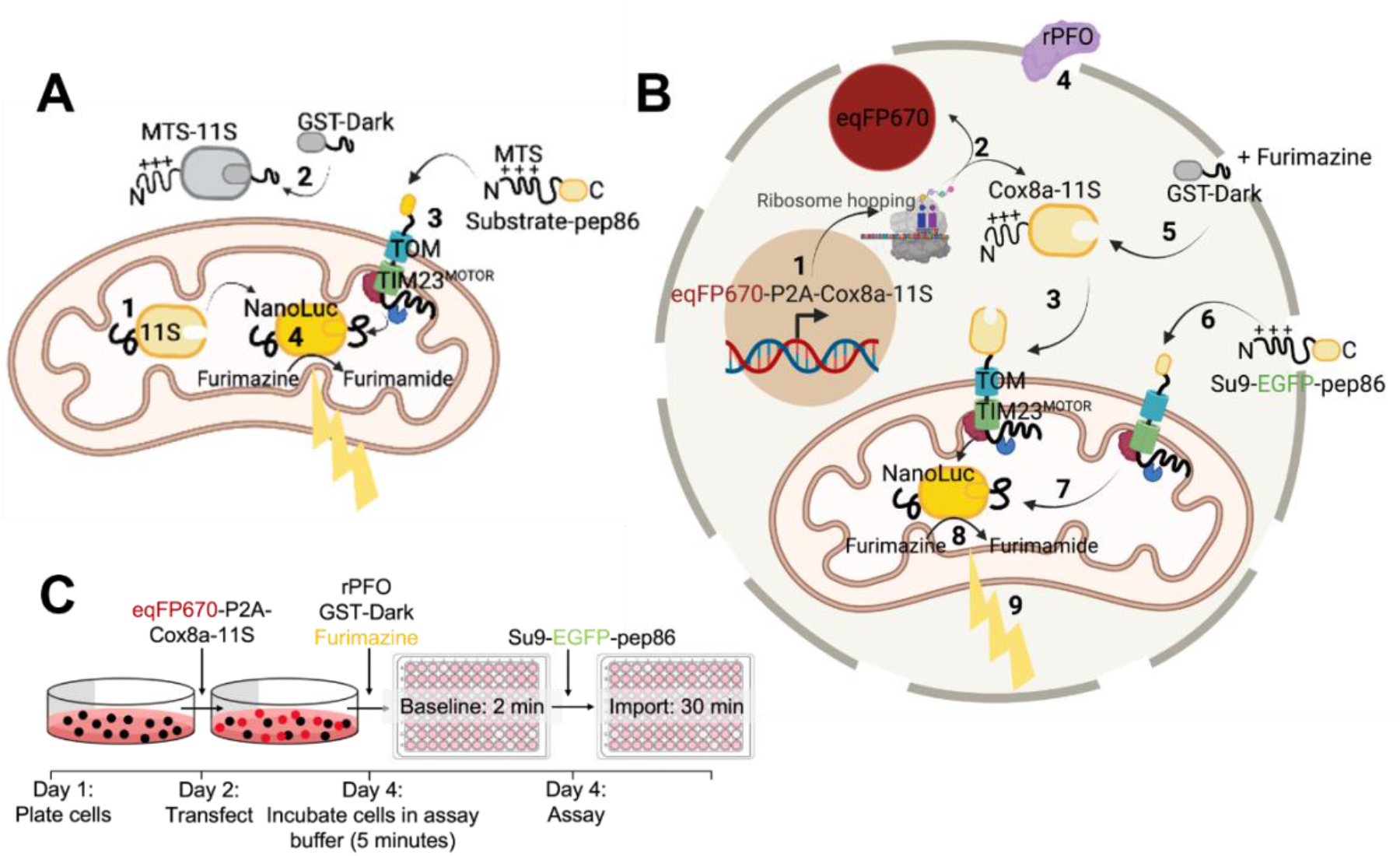
The NanoLuc Assay System for Monitoring Mitochondrial Protein Import in Yeast Mitochondria and Permeabilised Cells. **(A)** Schematic showing concept of the NanoLuc system to monitor mitochondrial import in isolated yeast mitochondria. Mitochondria were isolated from yeast producing a fusion of the mitochondrial targeting presequence (MTS) and the large fragment of split NanoLuc, 11S (MTS-11S). The resultant preparation with matrix localised 11S (with cleaved MTS) was deployed in the import assay: *(1)*. GST-Dark is added to bind non-mitochondrial 11S, decreasing background signal *(2)*. A pep86-tagged substrate protein (containing an N-terminal MTS; POI-pep86) is added and is targeted to the mitochondrial matrix *(3)*. The MTS is cleaved and pep86 binds to 11S in the matrix, forming NanoLuc *(4)*. This converts furimazine to furimamide producing a luminescent signal. Schematic created using BioRender. **(B)** Schematic showing the concept of the permeabilised NanoLuc import assay system in live mammalian cells. DNA encoding eqFP670 (far red fluorophore) followed by a P2A region and Cox8a-11S is transcribed in the nucleus and translated in the cytosol *(1)*, leading to the production of cytosolic eqFP670 and mitochondrially targeted Cox8a-11S *(2)*. Cox8a-11S is translocated to the mitochondrial matrix *via* the presequence pathway *(3)*. At the time of the assay, import buffer containing 3 nM rPFO is added, which perforates the plasma membrane *(4)* whilst retaining intact mitochondria. This allows other substrates, drugs, furimazine, and proteins to enter the cells. One such protein is GST-dark, which binds any remaining cytosolic 11S *(5)*, preventing background signal from cytosolic binding. Following a baseline read, a pep86 containing precursor is added (in this case Su9-EGFP-pep86; *(6)*) which is translocated into mitochondria where it binds to 11S *(7)*, forming the functional NanoLuc enzyme, which converts furimazine to furimamide *(8)*, producing a bioluminescent signal corresponding to import *(9)*. Schematic created using BioRender. **(C)** Experimental outline of the permeabilised NanoLuc assay. On day 1, cells are plated, and are subsequently transfected with eqFP670-P2A-Cox8a-11S DNA on day 2. On day 4, the day of the assay, media is replaced with assay buffer containing furimazine, rPFO, and GST-dark and incubated for 5 minutes prior to carrying out a 2-minute baseline read, followed by injection of Su9-EGFP-pep86 and a 30-minute kinetic import read for luminescence corresponding to protein import.

We first demonstrated the utility of this new system by monitoring protein import into mitochondria isolated from yeast [12]. In this case, 11S was constrained within the matrix for association with imported pep86 at the C-terminus of the precursor (precursor-pep86; Fig. 1A). The data was of sufficient quality to enable mathematical modelling approaches to help define the kinetic parameters underlying mitochondrial protein import in mitochondria [20]. Using this NanoLuc system we investigated how two major driving forces, ΔΨ and ATP hydrolysis, contribute to import through the TIM23^MOTOR^ complex, and how the process is affected by specific properties of the precursor protein [20]. These analyses could not have been achieved previously, particularly as the end point (amplitude) alone – the total amount of imported material of the assay, reported by the classic assay, is insufficient to understand the mechanism. Measuring both the amplitude and the rates led to a basic model for import, revealing a surprising and fundamental feature, whereby the reaction proceeds in two distinct steps: one at the outer membrane (TOM) and the other at the inner (TIM23).

While the new *in vitro* assay has proved insightful, it does have its limitations; particularly in view of the analysis of mitochondria in isolation. First of all, it appears that the recuperative powers of isolated mitochondria are insufficient to maintain the ΔΨ to drive multiple rounds of import through each translocation site [21]. Secondly, the interplay of the import machinery with regulatory cytosolic factors are all lost when studying purified mitochondria. This motivated us to adapt the assay to monitor import within intact cells. Such an assay would have the added advantage of observing fully fit mitochondria in their native state, that have not been subject to the physical trauma associated with isolation.

The assay we developed followed the principles devised for isolated yeast mitochondria (Fig. 1A) [12], but utilising permeabilised live mammalian cells (Fig. 1B). This in cellular version of the import assay provides accurate data with a high signal-to-noise ratio, comparable to the traces collected from isolated mitochondria, with the added bonus of context – *i.e*., the ability to relate import activity and failure to cell biology. The new in cell assay is outlined in Fig. 1B with a full experimental outline in Fig. 1C.

## Results

In live mammalian cells, the 11S component of the split luciferase was targeted to the matrix using the mitochondrial targeting presequence (MTS) of human Cox8a (a subunit of cytochrome *c* oxidase) (Fig. 1B). In order to quantify expression levels of 11S, which without the complementing peptide is colourless, a P2A bicistronic system was established to drive the stoichiometric production of the fluorescent protein eqFP670 [22]. This enabled the verification, quantification and normalisation of 11S expression, by virtue of the fluorescence co-expressed reporter: importantly, eqFP670 near-infrared excitation (Ex_eq670_ 605 nm) [23] is far enough away from 11S luminescence (405 nm) [11] to keep interference to a minimum.

### Characterisation and Optimisation of Cell Permeabilisation

Traditionally, cell permeabilisation for assessing mitochondrial physiology is accomplished using mild detergents such as digitonin or saponin. However, they are also known to permeabilise the mitochondrial outer membrane. Whilst this prospect is tolerable for assessing respiratory function, it is far from ideal for the analysis of protein import. So instead, we deployed recombinant perfringolysin (rPFO): a cholesterol-dependent, selective cytolysin [24], which has negligible mitochondrial side-effects up to 10 nM [25]. Upon oligomerisation, rPFO forms pores in the plasma membrane, allowing solutes and proteins of up to 200 kDa to rapidly equilibrate between the cytoplasm and extracellular medium [26, 27].

When cells are challenged with 5 mM succinate and 1 mM ADP, there is no respiratory response (change in oxygen uptake); this is expected, because they fail to permeate the plasma membrane (Fig. 2A). However, upon addition of 3 nM rPFO the rate of oxygen consumption increases rapidly (Fig. 2A), as succinate and ADP flood the cytosol and enter mitochondria through respective metabolite carrier proteins. Thus, demonstrating effective plasma membrane permeabilisation. The short delay prior to the burst of oxygen consumption is most likely due to the time taken for pore assembly. Importantly, the addition of exogenous cytochrome *c* during ADP-sustained respiration resulted in only a small change, indicating that the outer membrane was largely intact and had retained endogenous cytochrome *c*. The minor change elicited by the addition of cytochrome *c* nicely shows that exogenous proteins rapidly gain access to the mitochondrial surface. Therefore, rPFO treatment effectively permeabilises the plasma membrane to metabolites and proteins without compromising mitochondrial integrity or respiratory function.

**Fig. 2.**
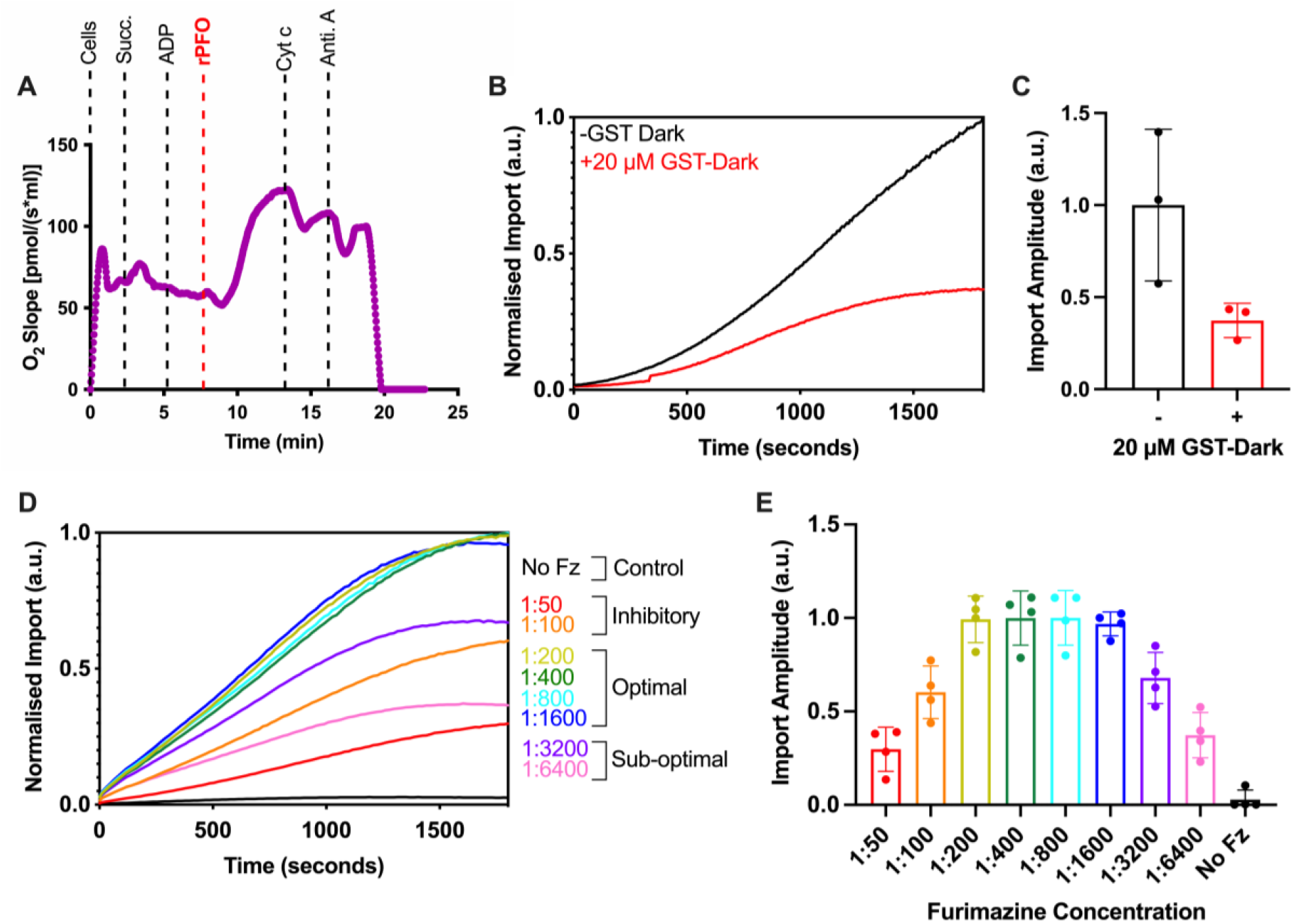
Permeabilised Cell NanoLuc Assay System Optimisation: rPFO, GST-Dark, and Fz. **(A)** Oroboros oxygraph trace showing mitochondrial respiration in response to various stimuli/poisons in HeLa cells. Cells are stimulated by addition of 5 mM succinate (Succ.) and 1 mM ADP prior to addition of 3 nM rPFO, which allows the substrates to access mitochondria. Cytochrome *c* (Cyt c; 10 μM) and antimycin A (Anti. A; 1 μM) are subsequently added, and the impact of these respiratory chain substrates on the oxygen consumption of the cells is measured. *Representative trace shown. N=3 biological replicates*. **(B)** Average traces from the NanoLuc import assay in the absence (black) or presence (red) of 20 μM GST-Dark, added to cells 5 minutes prior to the assay. Background was removed and data was normalised to cellular eqFP670 expression, and the maximum amplitude from the run, to allow comparison between runs regardless of raw values. *N=3 biological replicates, each with n=3 technical replicates*. **(C)** Maximum amplitude plotted from average import traces displayed in (B). *Error bars display SD*. **(D)** Average traces from the NanoLuc import assay showing import in the presence of various concentrations of furimazine (Fz; concentrations indicated as ratios of furimazine: assay buffer). Fz was applied to cells for 5 minutes prior to starting the assay, the NanoLuc assay was then carried out with a chasing precursor (Su9-EGFP-pep86). Data was normalised as described in (B). *N=4 biological replicates, each with n=3 technical replicates*. **(E)** Maximum amplitude plotted from average import traces displayed in (D). *Error bars display SD.*

A major issue with the NanoLuc import assay is the background signal arising from unimported 11S in the cytosol, release of internalised 11S or its presence in the external media resulting from lysed cells. To mitigate the background signal, we included an inactivated pep86 peptide fused to GST that can access extra-mitochondrial 11S, but not the matrix encapsulated reservoir [12]. A single amino acid substitution of a critical catalytic arginine residue of pep86 creates DarkBiT, which, upon binding to 11S, prevents catalysis by the formed NanoBiT complex [12, 28]. Thus, glutathione S-transferase fused to the DarkBiT peptide (GST-DarkBiT) quenches off-target luminescence arising from outside the matrix (Fig. 1A, B) [21].

Addition of purified GST-Dark to a final concentration of 20 μM (20-fold higher than precursor-pep86 fusion) considerably reduced the luminescence signal (Fig. 2B, C), which we attribute to binding and inactivation of residual 11S in the cytosol or external medium. This also prevents the non-mitochondrial 11S from sequestering precursor-pep86 *en route* to the mitochondria, and thereby ensuring the luminescent signal is solely produced by the formation of NanoLuc in the matrix [12, 21]. This quenching step is critical as evidently ~60% of the luminescence signal arises from non-matrix sources (Fig. 2B, C). Importantly, the remaining matrix signal is characteristic of import profiles seen with isolated mitochondria (Fig. 2B, red trace) [21].

At the same time as rPFO and GST-Dark addition to cells, the furimazine stock solution was added at a dilution of 1:800, as determined by a titration experiment to define the optimal dilution (Fig. 2D, E). At higher dilutions (over 1:200), furimazine has an inhibitory effect on mitochondrial import, probably due to respiratory toxicity, consistent with previous data in yeast mitochondria [12].

### The in-Cell NanoLuc Assay Requires ATP and PMF

As an import substrate, the engineered precursor protein Su9-EGFP-6xHis-pep86 was cloned for recombinant expression and purified in urea in an unfolded state. This protein contains the potent MTS of fungal Su9 for targeting to the mitochondrial matrix, as well as an EGFP fluorescent reporter. To characterise the specificity of the assay system for import *via* the presequence (TIM23^MOTOR^) pathway into the matrix, cells were treated with 1 μM antimycin A (A) – an inhibitor of complex III of the respiratory chain, and 5 μM Oligomycin (O) – an inhibitor of the ATP synthase. Together, these drugs limit the capacity to maintain ATP and PMF, both of which are required for protein import: PMF to drive the transport of positive residues across the membrane and ATP for the mitochondrial Hsp70 ATPase (mtHsp70) of the matrix located motor domain of the TIM23^MOTOR^ complex.

The impact of the AO inhibitors on respiratory function of the permeabilised cells was confirmed with a seahorse respirometer (Fig. 3A), corresponding to ~70% reduction in the amplitude (total amount of precursor import; Fig. 3B, C). Interestingly, compromising the available ATP and PMF affects the shape of the import curve (Fig. 3B, red trace): a reduction of a lag indicative of a profound impact on the kinetics of the import reaction [20]. The dependence of both the kinetics and total yield of import on ATP and PMF confirms that the luminescent signal is a *bona fide* measure of the mitochondrial presequence import pathway.

**Fig. 3.**
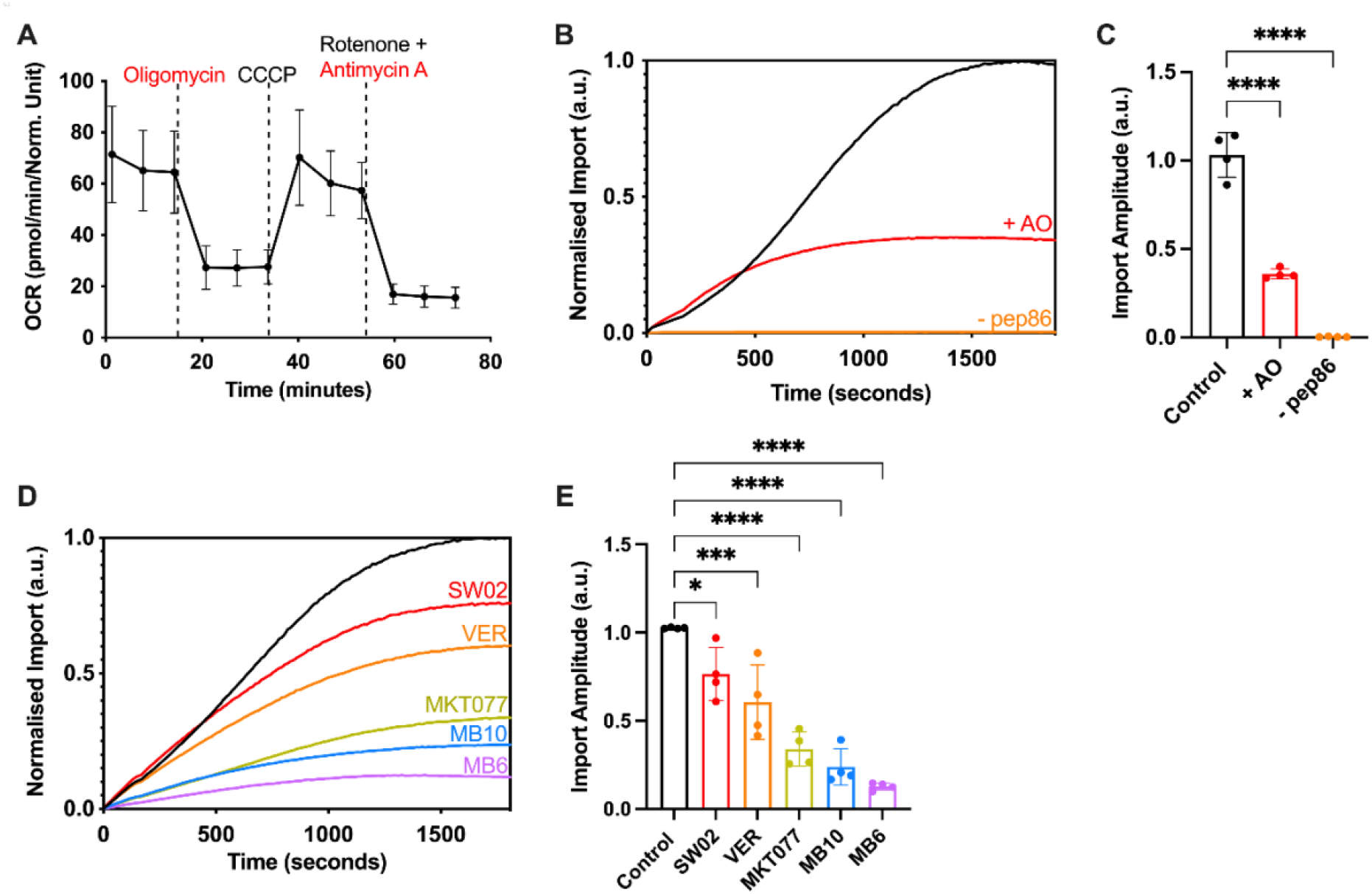
The Permeabilised NanoLuc Assay Monitors Import *via* the Presequence Pathway. **(A)** Mitochondrial stress test showing oxygen consumption rate (OCR) of cells when subjected to various respiratory chain substrates. HeLaGAL cells were grown for 48 hours prior to carrying out mitochondrial stress tests on a Seahorse XFe96. 1.5 μM oligomycin, 0.5 μM CCCP, 0.5 μM antimycin A, and 0.5 μM rotenone were added and their impact on mitochondrial oxygen consumption was measured. Data was normalised to cell density according to an SRB assay. *N=6 biological replicates, each with n=3 technical replicates*. **(B)** NanoLuc import trace in the absence (black) or presence (red) of PMF inhibitor AO (1 μM antimycin A, 5 μM oligomycin), or in the absence of pep86 containing precursor (orange). AO was applied to appropriate wells for 5 minutes prior to monitoring the import of the precursor (Su9-EGFP-pep86) using the NanoLuc assay. Resulting normalised average traces are shown. *N=4 biological replicates, each with n=3 technical replicates*. **(C)** Maximum amplitude plotted from average traces shown in (B). *Error bars display SD. One-way ANOVA and Tukey’s post hoc test were used*. **(D)** NanoLuc import traces in the presence of small molecule inhibitors of the import machinery. No drug (black), SW02 (20 μM, red), VER (20 μM, orange), MKT077 (20 μM, green), MB10 (30 μM, blue), or MB6 (10 μM, purple) were applied to cells for 5 minutes prior to starting the assay, the NanoLuc assay was then used to monitor import of a purified precursor (Su9-EGFP-pep86). Resulting normalised average traces are shown. *N=4 biological replicates, each with n=3 technical replicates*. **(E)** Maximum amplitude plotted from average traces shown in (D). *Error bars display SD. One-way ANOVA and Tukey’s post hoc test were used to determine significance*

Finally, the specificity of the assay was interrogated further by treating cells with drugs known to target various components of the import machinery (Fig. 3D, E). Cells were subject to an acute treatment by incubation of the drug 5 minutes prior to beginning the NanoLuc assays. Various small molecules were deployed which target the TIM23 presequence pathway: MB10, which inhibits mtHSP70 or TIM44 [10]; VER-155008, MKT077 and SW02 – all of which target the mtHSP70 ATPase [29–31]. MB6, an inhibitor of the MIA pathway [32] for protein targeting to the intermembrane space (IMS) [33].

All drug treatments were observed to induce a reduction in the total amount of imported precursor (Fig. 3D, E); unsurprisingly so, in the cases of VER-155008, MKT077, and MB10. Interestingly, SW02, which enhances the ATPase activity of mtHSP70 [31], increased the rate of import, but reduced the overall yield, suggestive of more subtle perturbation of the import process. The dramatic effect of MB6 on import efficiency was also surprising, considering that the imported substrate (Su9-EGFP-pep86) is not supposed to require the MIA pathway.

Whilst all the drugs have an impact on the amplitude, there are also variations in the rates and the shape and complexity of the curve. These are intriguing observations that warrant further investigation in the manner achieved for isolated yeast mitochondria [20]. Overall, the general dependencies and specific sensitivities we observed are consistent with faithful and accurate reporting of mitochondrial import.

## Discussion

We have developed, optimised, and road-tested an assay to monitor mitochondrial protein import in real-time with the capacity to obtain accurate kinetic data of events happening inside mammalian cells. The permeabilised cell NanoLuc assay works in HeLa cells and could be readily adapted for alternative immortalised or primary cell culture systems. Characterisation of the HeLa based system confirmed that the luminescence signal is an authentic measure of import *via* the presequence (TIM23^MOTOR^) pathway, due to its dependence on ATP and the PMF, and sensitivity to several specific and well characterised small molecule import inhibitors.

This system is based on the same concepts that we described for isolated yeast mitochondria [12, 21], enabled by permeabilisation using rPFO for accessibility of the necessary reagents, mitochondrial energisation, visualisation and experimentation (*e.g*., inhibition). Remarkably, the permeabilisation preserves mitochondrial respiratory capacity and, insofar as we can tell, has not significantly perturbed the system. Specifically, addition of cytochrome *c* showed that the mitochondrial outer membrane remained largely intact. Therefore, from the point of view of characterisation of protein import activity, regulation, and consequences of failure, the assay is robust and informative of mitochondrial physiology in the native cellular environment compared to an artificial stabilising solution. Importantly, the assay removes the need for mitochondrial isolation, and the inevitable physical damage this entails. Therefore, the NanoLuc assay has the capability to investigate import in an unperturbed state in fully functional mitochondria. Despite the clear advantages, we should also consider that freely diffusing cytosolic factors may be lost or diluted due to permeabilisation. So, it could be that some of the peripheral regulatory aspects of the import process may be impaired or lost, and not observable. Although, once identified lost regulatory factors could of course always be maintained exogenously. Future work will provide interesting and informative evaluation of how import in *in vivo* mitochondria compares to import into isolated mitochondria.

A key advance is that this new assay also provides opportunities to study the import process of mammalian mitochondria. Intact and functional mitochondria are challenging to isolate from cultured cells. So, avoiding this damaging step enables the analysis of a wide range of genetically manipulable mammalian cells. This is an important advance, freeing research of the restrictions of the classical model systems – beyond simple eukaryotes (yeast) and genetically intractable mammalian models (*e.g*., mitochondria isolated from cows and rats). Furthermore, this system has several benefits over classical import assays. As with the import assay into isolated mitochondria, the data acquisition time is reduced significantly from a matter of days to less than 30 minutes. Moreover, the assay can be run in a multiplate reader for rapid collection of large data sets at high time resolution. This high throughput capability will also facilitate small molecule screening ventures seeking drugs that modulate the import apparatus.

The permeabilised cell NanoLuc assay system can provide insight into the kinetics of import, aided by its sensitivity, allowing the detection of very small changes across all stages of the process. Furthermore, the high time resolution and quality of the data (low noise) enables mathematical fitting to kinetic models – as demonstrated for import into isolated yeast mitochondria [21] – for the dissection and characterisation of the different stages of import. Remarkably, the data presented here is of comparable quality, indicating it is now possible to do so in whole cell systems. Beyond this, we envisage that it will be possible to further develop this system to study the behaviour of particularly interesting import substrates – such as PINK1, the aberrant import of which has been linked to early onset Parkinson’s disease [34]. Likewise, we are now able to investigate further if and how certain aggregation prone protein variants (*e.g*., APP, Tau and Htt), associated with neurodegeneration, affect the import machinery.

The system is also adaptable to monitor alternative import pathways, *e.g., via* TIM23^SORT^ for lateral insertion into the IMM, MIA for IMS trafficking and the TIM22 complex for transport of presequence independent carrier family to the IMM. Incorporation of 11S in the IMS will also enrich our understanding of the various import routes, to visualise the transient entry of substrates to the TIM23 or TIM22 complexes, as well as the end point after IMS entry *via* MIA. Once the various import pathways can be monitored, the inhibitors tested here will help dissect the critical stages of each process.

A further goal will be to develop the assay system to primary cell cultures, such as neurons, and potentially also multi-cellular organisms. Aside from the additional challenges associated with the manipulation and imaging of primary cultures and model organisms, there are no technical barriers for the wider implementation of this assay into more interesting real-life situations. In this respect, it would be interesting to explore how mitochondrial import activity is affected in various models of disease, particularly those implicated with dysfunctional import.

## Materials & Methods

### Reagents

All chemicals were of the highest grade of purity commercially available and purchased from Sigma, UK, unless stated otherwise. Aqueous solutions were prepared in ultrapure water, while for non-aqueous solutions, ethanol or DMSO was used as solvent instead.

### Generation of Constructs

Constructs were generated by standard cloning techniques. PCR reactions were carried out using Q5 High Fidelity Hot Start DNA Polymerase (New England Biolabs (NEB), UK), using 20 pmol primers and 200 pg template DNA, as per manufacturers’ instructions. Restriction digest reactions were carried out using NEB restriction enzymes at 37°C for 45-60 min. Ligation reactions were carried out using T4 DNA Ligase (NEB) overnight at 16°C, using a ratio of 1:3 vector to insert (50 ng total). Transformation was carried out by incubation of 10% of total ligation mix/50 ng plasmid DNA construct in 50 μl competent *E. coli* cells (α-select, XL1-Blue, or BL21-DE3 cells were used, depending on application; all originally sourced from NEB and obtained from lab made stocks, stored at −80°C) for 30 min on ice, followed by heat shocking (45 sec, 42°C), and a further 15 min on ice. Cells were recovered by incubating in LB media at 37°C for 1 h and plated on LB-agar plates containing appropriate antibiotic. Plasmids were prepared by mini or maxi preps using commercially available kits (Qiagen, UK and Promega, UK, respectively) following manufacturers’ instructions, and verified by DNA sequencing using Eurofins Genomics, UK.

### Protein Purification

Protein expression was carried out exactly as described previously [12]. Pre-cultures of transformed BL21 (DE3) bacteria were grown overnight at 37°C in LB with appropriate antibiotic. Secondary cultures were inoculated at 1:100 from pre-cultures, in 1-8 L of 2X YT supplemented with appropriate antibiotic. Secondary cultures were grown until mid-log phase, then induced with 1 mM IPTG or 0.2% (w/v) Arabinose and grown for a further 3 h at 37°C. Cells were harvested by centrifugation, resuspended in ice-cold TK buffer (20 mM Tris, 50 mM KCl, pH 8.0), cracked in a cell disruptor at 4°C, and clarified by centrifugation at 38,000 rpm for 45 min at 4°C.

#### GST tagged Recombinant Perfringolysin (rPFO)

rPFO is a selective cytolysin [24], used as a permeabilisation agent in permeabilised cell NanoLuc import assays. Supernatant (soluble fraction) was loaded onto a 5 ml GSTrap 4B column (GE Healthcare, UK) and the column was washed with TK buffer. The peptide was eluted using freshly prepared 10 μM reduced glutathione in TK buffer. Eluted fractions were pooled and loaded onto a 5 ml anionic exchanger (Q-column; GE Healthcare). A salt gradient of 0-1 M KCl was applied at 5 ml/min over 20 min. The eluted fractions containing the protein were confirmed by SDS-PAGE with Coomassie staining, then pooled and concentrated by centrifugation at 4000 xg in a 50,000 MWCO PES VivaSpin concentrator (Sartorius) at 4°C. The final protein concentration was determined by A280 (extinction coefficient: 117,120 M^−1^ cm^−1^). The protein was aliquoted, snap frozen, and stored at −80°C. This protein was observed to have low stability once thawed, therefore a single aliquot was used per experiment.

#### GST-Dark Peptide

GST-Dark is an inactive version of pep86 (DarkBiT; [28]) fused to GST. It is used to quench any cytosolic 11S in import assays, preventing background signal from cytosolic binding of pep86 and 11S. GST-Dark was prepared as described previously [12]. Supernatant was loaded onto a GSTrap 4B column in TK buffer and purification was carried out exactly as for rPFO without the ion-exchange chromatography. Analysis, yield, and freezing was carried out exactly as for rPFO. For concentration, a 10,000 MWCO concentrator was used. Protein concentration was determined based on an extinction coefficient of 48,360 M^−1^ cm^−1^.

#### His tagged Su9-EGFP-pep86

Su9-EGFP-pep86 is a precursor protein with a Subunit 9 mitochondrial targeting pre-sequence (Su9-MTS) from *Neurospora crassa*, an EGFP reporter domain, and a pep86 peptide. Inclusion bodies (insoluble fraction) were solubilised in TK buffer supplemented with 6 M urea before loading onto a 5 ml HisTrap HP column (GE Healthcare). The column was first washed with TK + 6 M urea followed by TK + 6 M Urea + 50 mM imidazole. The protein was eluted in 300 mM imidazole. Eluted fractions containing the desired protein were pooled and loaded onto a 5 ml cationic exchanger (S-column; GE Healthcare). A salt gradient of 0-1 M KCl was applied over 20 min (at 5 ml/min) and the protein was eluted in 5 ml fractions. Imidazole was removed by spin concentration, followed by dilution in TK buffer containing 6 M urea. Analysis, yield, and freezing were carried out exactly as for rPFO. For concentration, a 10,000 MWCO concentrator was used. Protein concentration was determined based on an extinction coefficient of 28,880 M^−1^ cm^−1^.

### Cell Culture

HEK293T (ECACC) and glucose-grown HeLa (HeLaGLU; ATCC) cells were maintained in Dulbecco’s Modified Eagle’s Medium (DMEM; Gibco, UK) supplemented with 10% (v/v) foetal bovine calf serum (FBS; Invitrogen, UK) and 1% (v/v) penicillin-streptomycin (P/S; Invitrogen). Where OXPHOS dependence was required, HeLa cells were cultured in galactose medium (HeLaGAL) consisting of DMEM without glucose (Gibco) supplemented with 10 mM galactose, 1 mM sodium pyruvate, 4 mM glutamine, 10% FBS and 1% P/S. Cells were cultured in galactose media for at least 3 weeks prior to experiments on HeLaGAL cells. Cells were maintained in T75 ventilated flasks in humidified incubators at 37°C with 5% CO_2_.

### Cell Transduction by Transfection

HeLa cells were seeded and grown up to 70-80% confluency. At this point cells were transfected with 0.5-2 μg of the desired DNA using Lipofectamine 3000 reagent (Thermo Fisher Scientific, UK) using a 1:3 DNA to lipofectamine ratio and following the manufacturers’ protocol. Cells were then grown for a further 24-72 hours prior to experimental analysis.

### Cell Transduction by Lentiviral Infection

Lentiviral particles were produced in HEK293T cells by addition of a mixture of DNAs (27.2 μg DNA to be produced, and packaging vectors pMDG2 (6.8 μg) and pAX2 (20.4 μg)) and pEI transfection reagent (1.5:1 pEI:DNA to be produced) in OptiMEM media (Gibco) to cells in a T75 flask at 70-80% confluency. After 6 h, media was changed to complete DMEM, and cells incubated for 72 h to allow lentivirus particle production. Lentivirus particles were harvested from the media at 48 and 72 h for maximum yield, pooled, spun down at 4000 xg for 5 min to remove dead cells, and concentrated by adding Lenti-X concentrator (Takara Bio) at a 1:3 ratio and incubating at 4°C for at least 1 h. Lentiviral particles were pelleted by centrifugation at 4000 xg for 45 min. Pellets were resuspended in plain DMEM at 1:50 of initial supernatant volume, aliquoted, and stored at −80°C until required. For infection, 5-20 μl concentrated lentivirus (per 35mm dish, scaled up accordingly and optimised by titration of each fresh batch) was added dropwise to cell media when cells reached 70-80% confluency. Cells were incubated with lentivirus for 24-72 h prior to experimental analysis.

### NanoLuc Assay in Permeabilised Cells

Cells were seeded in 6-well plates (300,000 cells/well), transfected with eqFP670-P2A-Cox8a-11S, and incubated for 48 h at 37°C (with additional lentiviral transduction carried out 24 h after 11S transfection, where appropriate). After 48 h of 11S transfection, cells were seeded on standard white flat-bottom 96-well plates at 30,000 cells/well and import assays performed 16 h afterwards. Then, cells were washed three times with 200 μl HBSS (calcium and magnesium free) and incubated in HBSS imaging buffer (1 X HBSS (Thermo), 5 mM D-(+)-glucose,10 mM HEPES (Santa Cruz Biotechnology, Germany), 1 mM MgCl_2_, 1 mM CaCl_2_, pH 7.4). A fluorescence read was taken at 605/670nm using monochromators with gain set to allow maximum sensitivity without saturation, using a BioTek Synergy Neo2 plate reader, for normalisation of results against eqFP670 expression. Buffer was then removed and cells were incubated in permeabilised cell assay master mix (225 mM mannitol, 10 mM HEPES, 2.5 mM MgCl_2_, 40 mM KCl, 2.5 mM KH_2_PO_4_, 0.5 mM EGTA, pH 7.4) supplemented with 5 mM succinate, 1 μM rotenone, 0.1 mg/ml creatine kinase, 5 mM creatine phosphate, 1 mM ATP, 0.1% (v/v) Prionex, 3 nM rPFO (purified in house), 20 μM GST-Dark (purified in house), 1:800 furimazine (Nano-Glo® Luciferase Assay System; Promega). Drugs or inhibitors were added to individual wells as described in figure legends. A baseline read of 30 sec of background luminescent signal was taken prior to injection of purified substrate protein (Su9-EGFP-pep86) to 1 μM final concentration, followed by a further bioluminescence read corresponding to import, lasting 30 min. Bioluminescence was read using a BioTek Synergy Neo2 plate reader (Agilent, UK) or a CLARIOstar Plus plate reader (BMG LabTech, UK) without emission filters with gain set to allow maximum sensitivity without saturation, and with acquisition time of 0.1 sec per well. Row mode was used, and reads were taken every 6 sec or less, with wells in triplicate.

### Mitochondrial Respiration Analysis

#### Oxygraph Assay

Cells were grown to confluency in T75 flasks, then harvested with trypsin, pelleted and washed once with HBSS (Ca^2+^- and Mg^2+^-free), and finally resuspended in mannitol respiration buffer (225 mM mannitol, 10 mM HEPES, 2.5 mM MgCl_2_, 40 mM KCl, 2.5 mM KH_2_PO_4_, 0.5 mM EGTA, pH 7.4). Respiratory function assays were carried out in a Oxygraph-2k (Oroboros Instruments, Innsbruck, Austria) at 37 °C in a 2 ml closed chamber with 5 million cells per run. Drugs were added as indicated in figure legends and respiration rates were calculated as the average value over a 45 sec window in DatLab 5 (Oroboros Instruments) and are expressed in nmol O_2_ / min /mg protein.

#### Seahorse Assay: Mitochondrial Stress Test

Cells were seeded in 6-well plates (300,000 cells/well) and infected with the appropriate lentivirus, as described in figure legends. The day prior to the assay, cells were seeded at 10,000 cells/well in 96 well Seahorse XF cell culture plates (Agilent, UK). The sensor cartridges were hydrated overnight with tissue culture grade H_2_O in a non-CO_2_ incubator at 37°C as per manufacturer’s instructions. On the day of the assay, H_2_O in the sensor plate was replaced with Seahorse XF Calibrant (Agilent) and cells were washed with HBSS and incubated in Seahorse media (Seahorse XF assay medium (Agilent), 1 mM pyruvate (Agilent), 2 mM glutamine (Agilent), 10 mM D-(+)-galactose). Both sensor and cell plates were incubated in a non-CO_2_ incubator at 37°C for 1 h prior to the assay. The sensor plate was loaded with oligomycin (15 μM for 1.5 μM final concentration in wells; injector A), CCCP (5 μM for 0.5 μM final; injector B) and antimycin A and rotenone (5 μM/5 μM for 0.5 μM/0.5 μM final; injector C). The sensor plate was calibrated in the machine prior to loading cells and running a mitochondrial stress test using a Seahorse XFe96 analyser (Agilent). Following assays, cells were washed and fixed in 1% acetic acid in methanol at −20°C overnight for SRB assays, which were used for normalising data to protein content.

### SRB Assay

For analysis of cell density based on cellular protein content, the sulforhodamine B (SRB) assay was used [369]. Cells were fixed with ice cold 1% (v/v) acetic acid in methanol overnight at −20°C. Fixative was aspirated and plates were allowed to dry at 37°C before proceeding to the next step. SRB (0.5% (w/v) in 1% (v/v) acetic acid in dH2O) was incubated in wells (37°C, 30 minutes). SRB was then aspirated, and unbound stain removed by cells extensive washing using 1% acetic acid, prior to drying plates at 37°C. Bound protein stain was solubilised by shaking incubation with 10 mM Tris (pH 10; 15 min, RT). Absorbance was read on a microplate reader with a 544/15 nm filter.

### Statistical Analysis

Statistical significance between groups was determined using unpaired Student’s t-Test or one and two-way ANOVA with interaction if more than two groups were analysed. Following ANOVA, p values were adjusted for multiple comparisons through Tukey’s or Dunnett’s post-hoc tests and differences were considered significant at 5% level. Statistical analyses were performed using Graph Pad Prism version 9 (GraphPad Software, Inc., San Diego, CA, USA).

## Author contribution

HIN, GCP and IC designed experiments; HIN conducted all experiments; HIN and IC wrote the manuscript; GCP and JMH edited the manuscript; IC secured funding and led the project.

## Declaration

The authors declare no competing interests.

## Data and materials availability

All data are available in the main text or the supplementary materials.

## References

[1] Neupert W. Protein import into mitochondria. Annual Review of Biochemistry. 1997;66:863–917.

[2] Needs HI, Protasoni M, Henley JM, Prudent J, Collinson I, Pereira GC. Interplay between Mitochondrial Protein Import and Respiratory Complexes Assembly in Neuronal Health and Degeneration. Life-Basel. 2021;11.

[3] Sokol AM, Sztolsztener ME, Wasilewski M, Heinz E, Chacinska A. Mitochondrial protein translocases for survival and wellbeing. Febs Letters. 2014;588:2484–95.

[4] Devi L, Prabhu BM, Galati DF, Avadhani NG, Anandatheerthavarada HK. Accumulation of amyloid precursor protein in the mitochondrial import channels of human Alzheimer’s disease brain is associated with mitochondrial dysfunction. Journal of Neuroscience. 2006;26:9057–68.

[5] Devi L, Raghavendran V, Prabhu BM, Avadhani NG, Anandatheerthavarada HK. Mitochondrial import and accumulation of alpha-synuclein impair complex I in human dopaminergic neuronal cultures and Parkinson disease brain. Journal of Biological Chemistry. 2008;283:9089–100.

[6] Yano H, Baranov SV, Baranova OV, Kim J, Pan YC, Yablonska S, et al. Inhibition of mitochondrial protein import by mutant huntingtin. Nature Neuroscience. 2014;17:822–31.

[7] Di Maio R, Barrett PJ, Hoffman EK, Barrett CW, Zharikov A, Borah A, et al. alpha-Synuclein binds to TOM20 and inhibits mitochondrial protein import in Parkinson’s disease. Science Translational Medicine. 2016;8:14.

[8] Kang PJ, Ostermann J, Shilling J, Neupert W, Craig EA, Pfanner N. REQUIREMENT FOR HSP70 IN THE MITOCHONDRIAL MATRIX FOR TRANSLOCATION AND FOLDING OF PRECURSOR PROTEINS. Nature. 1990;348:137–43.

[9] Miyata N, Steffen J, Johnson ME, Fargue S, Danpure CJ, Koehler CM. Pharmacologic rescue of an enzyme-trafficking defect in primary hyperoxaluria 1. Proceedings of the National Academy of Sciences of the United States of America. 2014;111:14406–11.

[10] Miyata N, Tang ZY, Conti MA, Johnson ME, Douglas CJ, Hasson SA, et al. Adaptation of a Genetic Screen Reveals an Inhibitor for Mitochondrial Protein Import Component Tim44. Journal of Biological Chemistry. 2017;292:5429–42.

[11] Dixon AS, Schwinn MK, Hall MP, Zimmerman K, Otto P, Lubben TH, et al. NanoLuc Complementation Reporter Optimized for Accurate Measurement of Protein Interactions in Cells. Acs Chemical Biology. 2016;11:400–8.

[12] Pereira GC, Allen WJ, Watkins DW, Buddrus L, Noone D, Liu X, et al. A High-Resolution Luminescent Assay for Rapid and Continuous Monitoring of Protein Translocation across Biological Membranes. Journal of Molecular Biology. 2019;431:1689–99.

[13] Smoyer CJ, Katta SS, Gardner JM, Stoltz L, McCroskey S, Bradford WD, et al. Analysis of membrane proteins localizing to the inner nuclear envelope in living cells. Journal of Cell Biology. 2016;215:575–90.

[14] Wehrman TS, Casipit CL, Gewertz NM, Blau HM. Enzymatic detection of protein translocation. Nature Methods. 2005;2:521–7.

[15] Krayl M, Guiard B, Paal K, Voos W. Fluorescence-mediated analysis of mitochondrial preprotein import in vitro. Analytical Biochemistry. 2006;355:81–9.

[16] Hall MP, Unch J, Binkowski BF, Valley MP, Butler BL, Wood MG, et al. Engineered Luciferase Reporter from a Deep Sea Shrimp Utilizing a Novel Imidazopyrazinone Substrate. Acs Chemical Biology. 2012;7:1848–57.

[17] England CG, Ehlerding EB, Cai WB. NanoLuc: A Small Luciferase Is Brightening Up the Field of Bioluminescence. Bioconjugate Chemistry. 2016;27:1175–87.

[18] Allen WJ, Watkins DW, Dillingham MS, Collinson I. Refined measurement of SecA-driven protein secretion reveals that translocation is indirectly coupled to ATP turnover. Proceedings of the National Academy of Sciences of the United States of America. 2020;117:31808–16.

[19] Allen WJ, Corey RA, Watkins DW, Oliveira ASF, Hards K, Cook GM, et al. Rate-limiting transport of positively charged arginine residues through the Sec-machinery is integral to the mechanism of protein secretion. Elife. 2022;11:23.

[20] Ford HC, Allen WJ, Pereira GC, Liu X, Dillingham MS, Collinson I. Towards a molecular mechanism underlying mitochondrial protein import through the TOM and TIM23 complexes. Elife. 2022;11.

[21] Ford HC, Allen WJ, Pereira GC, Liu X, Dillingham MS, Collinson I. Towards a molecular mechanism underlying mitochondrial protein import through the TOM and TIM23 complexes. bioRxiv. 2021:2021.08.30.458282.

[22] Daniels RW, Rossano AJ, Macleod GT, Ganetzky B. Expression of Multiple Transgenes from a Single Construct Using Viral 2A Peptides in Drosophila. Plos One. 2014;9:10.

[23] Shcherbo D, Shemiakina, II, Ryabova AV, Luker KE, Schmidt BT, Souslova EA, et al. Near-infrared fluorescent proteins. Nature Methods. 2010;7:827–U1520.

[24] Kacprzyk-Stokowiec A, Kulma M, Traczyk G, Kwiatkowska K, Sobota A, Dadlez M. Crucial Role of Perfringolysin O D1 Domain in Orchestrating Structural Transitions Leading to Membrane-perforating Pores A HYDROGEN-DEUTERIUM EXCHANGE STUDY. Journal of Biological Chemistry. 2014;289:28738–52.

[25] Divakaruni AS, Wiley SE, Rogers GW, Andreyev AY, Petrosyan S, Loviscach M, et al. Thiazolidinediones are acute, specific inhibitors of the mitochondrial pyruvate carrier. Proceedings of the National Academy of Sciences of the United States of America. 2013;110:5422–7.

[26] Rossjohn J, Polekhina G, Feil SC, Morton CJ, Tweten RK, Parker MW. Structures of perfringolysin O suggest a pathway for activation of cholesterol-dependent cytolysins. Journal of Molecular Biology. 2007;367:1227–36.

[27] Ramachandran R, Heuck AP, Tweten RK, Johnson AE. Structural insights into the membrane-anchoring mechanism of a cholesterol-dependent cytolysin. Nature Structural Biology. 2002;9:823–7.

[28] Dixon AS, Encell L, Hall M, Wood K, Wood M, Schwinn M, et al. Activation of bioluminescence by structural complementation. USA: Promega Corporation; 2017.

[29] Williamson DS, Borgognoni J, Clay A, Daniels Z, Dokurno P, Drysdale MJ, et al. Novel Adenosine-Derived Inhibitors of 70 kDa Heat Shock Protein, Discovered Through Structure-Based Design. Journal of Medicinal Chemistry. 2009;52:1510–3.

[30] ModicaNapolitano JS, Koya K, Weisberg E, Brunelli BT, Li Y, Chen LB. Selective damage to carcinoma mitochondria by the rhodacyanine MKT-077. Cancer Research. 1996;56:544–50.

[31] Wisen S, Bertelsen EB, Thompson AD, Patury S, Ung P, Chang L, et al. Binding of a Small Molecule at a Protein-Protein Interface Regulates the Chaperone Activity of Hsp70-Hsp40. Acs Chemical Biology. 2010;5:611–22.

[32] Dabir DV, Hasson SA, Setoguchi K, Johnson ME, Wongkongkathep P, Douglas CJ, et al. A Small Molecule Inhibitor of Redox-Regulated Protein Translocation into Mitochondria. Developmental Cell. 2013;25:81–92.

[33] Stojanovski D, Muller JM, Milenkovic D, Guiard B, Pfanner N, Chacinska A. The MIA system for protein import into the mitochondrial intermembrane space. Biochimica Et Biophysica Acta-Molecular Cell Research. 2008;1783:610–7.

[34] Pickrell AM, Youle RJ. The Roles of PINK1, Parkin, and Mitochondrial Fidelity in Parkinson’s Disease. Neuron. 2015;85:257–73.

